# mBeRFP, a versatile fluorescent tool to enhance multichannel live imaging and its applications

**DOI:** 10.1101/2022.01.03.474820

**Authors:** Emmanuel Martin, Magali Suzanne

## Abstract

Cell and developmental biology increasingly require live imaging of protein dynamics in cells, tissues or living organisms. Thanks to the discovery and the development of a panel of fluorescent proteins over the last decades, live imaging has become a powerful and commonly used approach. However, multicolor live imaging remains challenging. The generation of long Stokes shift red fluorescent proteins, such as mBeRFP, offers interesting new perspectives to bypass this limitation. Here, we constructed a set of mBeRFP-expressing vectors and provided a detailed characterization of this fluorescent protein for *in vivo* live imaging and its applications in *Drosophila*. Briefly, we showed that a single illumination source is sufficient to simultaneously stimulate mBeRFP and GFP. We demonstrated that mBeRFP can be easily combined with classical green and red fluorescent protein without any crosstalk. We also showed that the low photobleaching of mBeRFP is suitable for live imaging, and that this protein can be used for quantitative applications such as FRAP or laser ablation. Finally, we believe that this fluorescent protein, with the set of new possibilities it offers, constitutes an important tool for cell, developmental and mechano biologists in their current research.

## Introduction

The field of cell and developmental biology relies more and more on the analysis of protein dynamics in cells, tissues or organisms using live imaging. In addition to major advances in fluorescence microscopy, the discovery of the Green Fluorescent Protein (GFP) in the jellyfish *Aequorea victoria* (SHIMOMURA et al., 1962) and its first use as a fluorescent marker in non-jellyfish organisms (Chalfie et al., 1994) were a critical breakthrough for the field of non-invasive protein imaging. Since GFP discovery, many other fluorescent proteins (FPs) have been identified, developed and enhanced, providing a powerful toolkit for visualization of dynamic processes *in vivo* (Chudakov et al., 2010). However, in classical fluorescence imaging, i.e. without a spectral detector and linear unmixing, the number of fluorescent proteins that can be tracked in a living tissue is often limited by the intrinsic spectral properties of FPs to avoid crosstalk as well as by the instrumentation available. Indeed, to detect specifically two FPs simultaneously, their excitation and emission peak should be separated by at least 60 nm (Thorn, 2017); so usually only the pairs CFP / YFP, or the commonly used GFP / RFP are imaged together. In some cases, with an adapted set of filters, a third color can be added with little crosstalk such as RFP with the couple CFP / YFP (Boulina et al., 2013), or BFP (mTagBFP2 (Subach et al., 2011)) with GFP / RFP. However, in this latter case, a near-UV excitation is required, which is highly phototoxic for long-term live imaging experiments (Gorgidze et al., 1998; Icha et al., 2017). Thus, combining more than two colors for live imaging with low phototoxicity, sufficient brightness and photostability of FPs appears challenging, especially in model organisms where the environment completely differs from assessments done *in vitro* to characterize properties of these FPs (Heppert et al., 2016; Wiedenmann et al., 2009).

During the last two decades, an interest in monomeric red fluorescent proteins characterized by a large Stokes shift (LSS RFP, i.e., a difference between the emission peak and the excitation peak larger than 150 nm) appeared. These proteins are particularly interesting because they can be excited by blue light allowing the simultaneous excitation of the LSS RFP with CFP or GFP. Among these LSS RFP, mKeima (Kogure et al., 2006), LSSmKate1, LSSmKate2 (Piatkevich et al., 2010a; Piatkevich et al., 2010b) and mBeRFP (Yang et al., 2013) were generated. However, their full potential in living organisms has not been tested yet.

Here, we provided a detailed characterization of mBeRFP for *in vivo* live imaging applications in *Drosophila*. First, we cloned and expressed in flies two different LSS RFP, LSSmKate2 and mBeRFP, and evaluated their possible use in live imaging. Because of its higher fluorescence intensity, we focused on mBeRFP and showed that it can be easily combined with green and classical red fluorescent proteins or dyes without crosstalk. Finally, we demonstrated that mBeRFP can be combined to eGFP in live imaging, offering the interesting possibility to image the two fluorophores using a single light source. We also showed the potential interest of this protein for quantitative and biophysical applications such as FRAP or laser ablation.

## Results and Discussion

### A set of mBeRFP-expressing vectors

To test the potential of fluorescent extended Stokes shift proteins in *Drosophila*, we constructed a set of plasmids expressing either cytoplasmic or nuclear mBeRFP under the control of a UAS promoter (5xUAS + hsp70 minimal promoter - Fig.1A), as well as a cytoplasmic LSSmKate2, and generated the corresponding transgenic flies. To ensure that the expression of mBeRFP or LSSmKate2 did not induce developmental defects or cytotoxicity, we expressed these constructs in leg and wing discs using different Gal4 promoters (Fig.1B and C; FigSup1A). This revealed that morphogenesis was not affected by the expression of both cytoplasmic and nuclear mBeRFP or LSSmKate2. However, the fluorescence intensity of the LSSmKate2 was not as high as the one of mBeRFP (FigSup1B), so we focused on mBeRFP.

**Figure 1:**
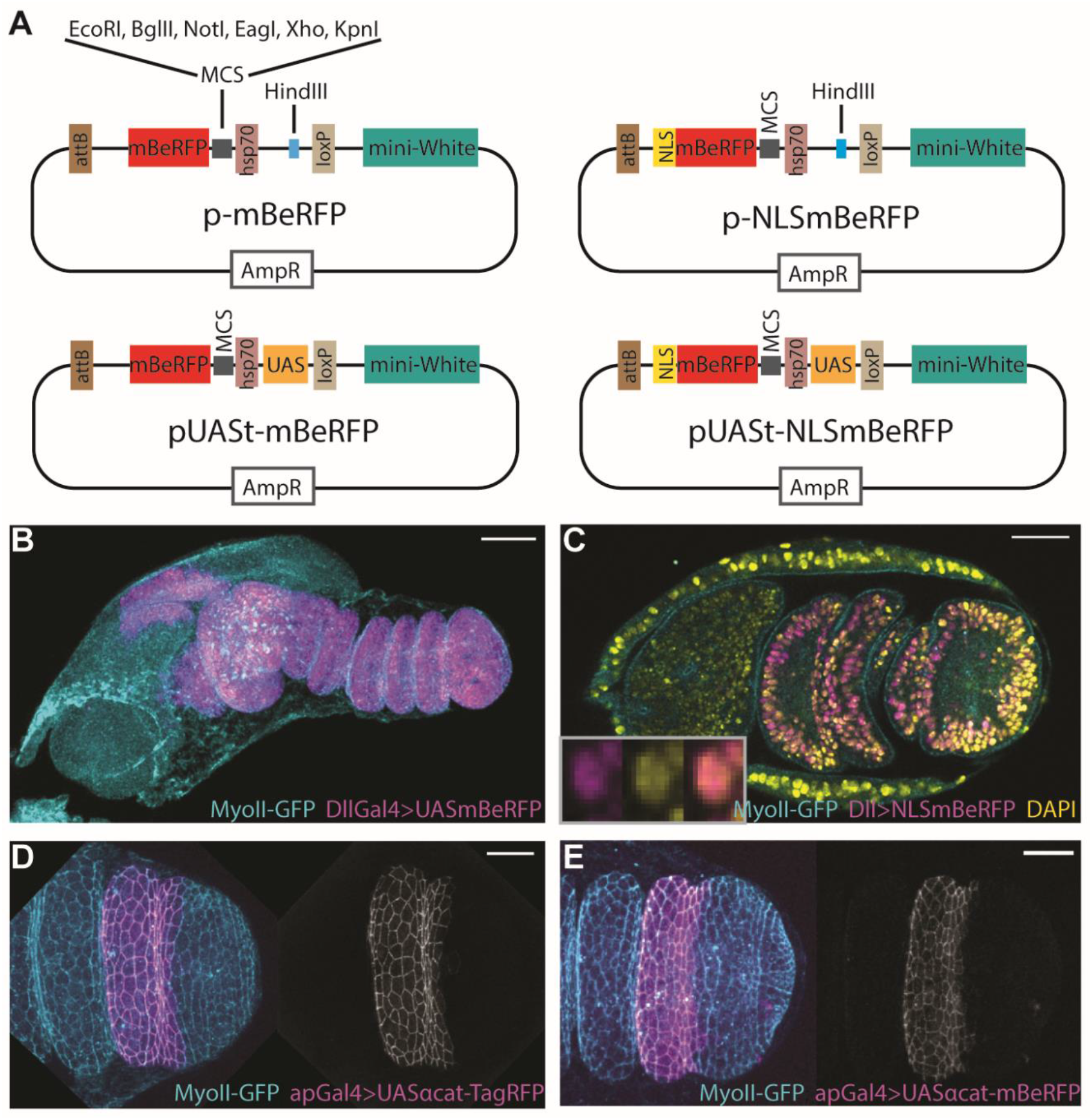
A set of mBeRFP-expressing vectors for *Drosophila*. **A)** The key features of mBeRFP vectors. The restrictions sites available in the MCS are listed only once but are the same for all constructs. The hsp70 minimal promoter is not schematized even if present in all vectors. **B)** Z-projection of an imaginal leg disc expressing endogenous MyoII-GFP (sqh-eGFP, cyan) and cytoplasmic mBeRFP (magenta) in the Dll domain of expression. **C)** Cross-section of leg epithelium showing the expression of nuclear mBeRFP (magenta) in the Dll expression domain and MyoII-GFP (cyan), stained with DAPI (yellow). **D-E)** Distal part of leg epithelium expressing endogenous MyoII-eGFP (cyan) and UAS alpha-catenin-TagRFP (D, magenta or grey) or UAS alpha-catenin-mBeRFP (E, magenta or grey) in the apterous expression domain. Scale bar represents 40 µm in B, C and 20 µm in D, E.

We then tested the possibility to use mBeRFP as a tag to follow endogenous protein dynamics. As a proof of concept, we inserted the alpha-catenin coding sequence in frame with the mBeRFP in the pUASt-mBeRFP plasmid and expressed the resulting α-catenin-mBeRFP fusion protein in a restricted domain of the leg epithelium (*apterous* domain). We used α-catenin-TagRFP-expressing flies (Gal4/UAS driven system) as a control. No significant difference in terms of location and morphogenesis was observed between these two fusion proteins (Fig.1 D and E) suggesting that mBeRFP, alone or fused to proteins of interest, could be used *in vivo* in *Drosophila*. In addition, we developed a ‘promoter-free’ version (containing only the hsp70 minimal promoter), allowing versatile applications such as the expression of cytoplasmic, nuclear or fused version of mBeRFP after integration of a promoter of interest at the HindIII restriction site (Fig.1A).

### mBeRFP can be combined with green and red FPs without crosstalk

One of the interests of using new fluorescent proteins is the possibility to combine it with commonly used fluorescent proteins, such as GFP or RFP. Thus, to ensure that the mBeRFP protein can be used in combination with these classical fluorescent proteins, we observed their respective emission in function of the excitation wavelength and analyzed the crosstalk. To test the combination with green-emitting fluorescent proteins, we used enhanced Green Fluorescent Protein (eGFP). Theoretically, eGFP and mBeRFP can be coupled without crosstalk using adapted emission windows (Fig.2A). In practice, we fixed emission windows from 490 nm to 550 nm for eGFP and from 600 nm to 700 nm for mBeRFP (Fig.2A). While the maximal excitation wavelength differs between these two proteins - 488 nm and 446 nm for eGFP and mBeRFP respectively - the higher brightness of eGFP compared to mBeRFP (Thorn, 2017) offers the interesting possibility to excite the two proteins with the same wavelength. Indeed, using the excitation wavelength at 458 nm, we could excite mBeRFP to more than 90% and eGFP to 60% (Fig.2A). We imaged at the same time alpha-catenin-mBeRFP expressed under the control of an apterous-Gal4 promoter and the endogenous eGFP-tagged Myosin II using either a classical confocal mode or a spectral mode to completely avoid potential spectral fluorescence overlap and crosstalk (Fig.2B)(Zimmermann et al., 2003). We calculated the crosstalk in classical laser scanning mode as the difference between the rate of eGFP emitted in the mBeRFP emission window when imaged with the spectral mode and the corresponding rate when imaged with the classical mode. This measure revealed the total absence of crosstalk between eGFP and mBeRFP with these settings (Fig.2C) confirming the possible combination of these two proteins. To further test the specificity and the precision in terms of spatial distribution of the two fusion proteins eGFP and mBeRFP using a single excitation wavelength, we used random illumination microscopy, a newly developed super-resolution technique (Mangeat et al., 2021). We concomitantly imaged MyoII-GFP and alpha-catenin-mBeRFP with a single illumination source at 445 nm and were able to resolve distinctly the two distinct cortical myosin belts of neighboring cells separated by the adherens junctions, without fluorescence crosstalk (FigSup.2A and B).

**Figure2:**
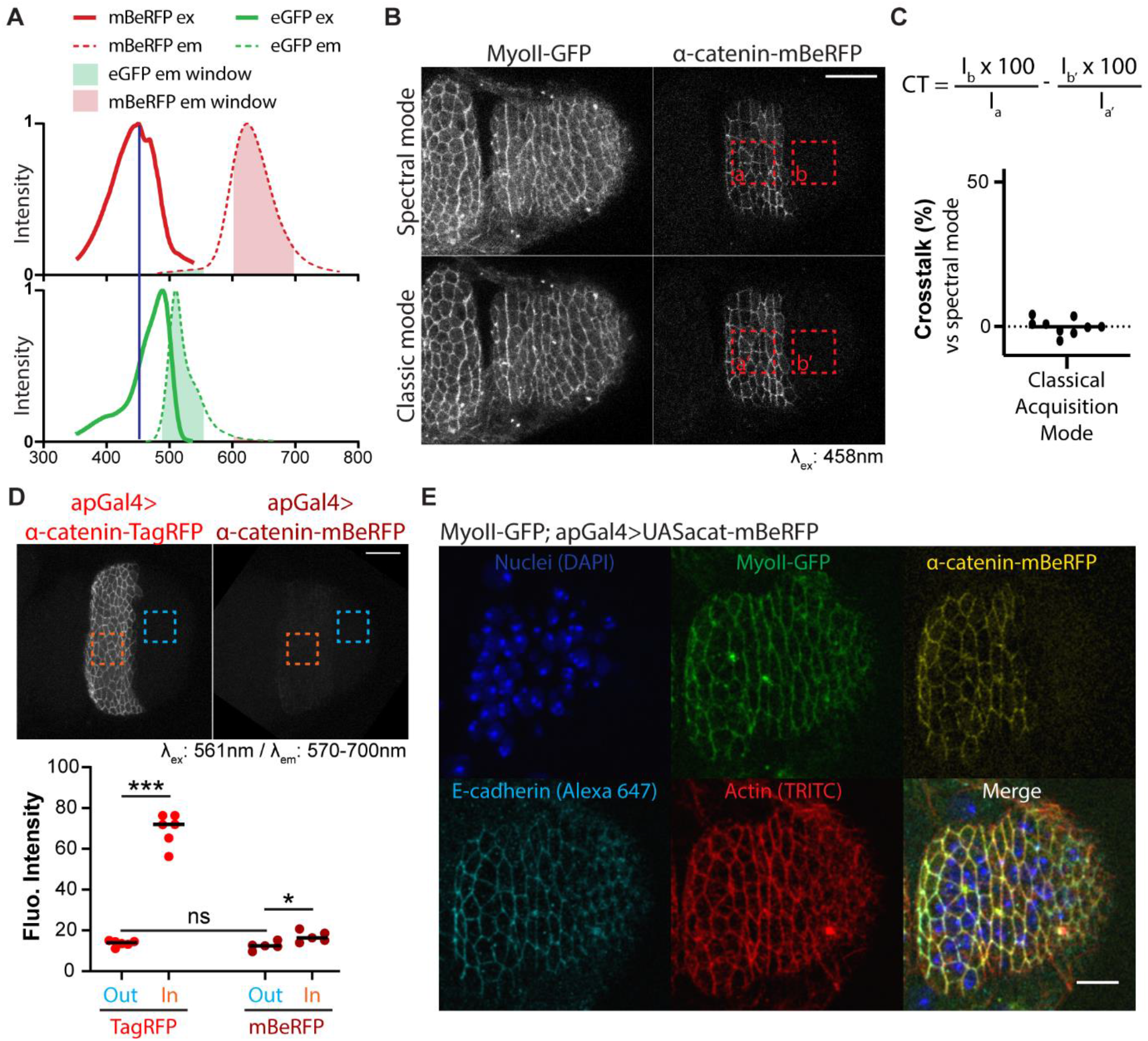
mBeRFP can be combined with green- and red-emitting fluorescent proteins. **A)** mBeRFP and eGFP excitation and emission spectra showing eGFP and mBeRFP emission windows (respectively 490 to 550 nm and 600 to 700 nm) as well as excitation wavelength (dark blue line - 458 nm) used. **B-C)** Crosstalk between eGFP and mBeRFP. **B)** Z-projection of the distal part of a leg disc showing MyoII-GFP (sqh-eGFP) and UAS alpha-catenin-mBeRFP imaged using spectral (top) or classical confocal mode (bottom). Dashed squares point out areas used to calculate the crosstalk in C. **C)** Dot plot representing the crosstalk (CT) together with the corresponding formula. **D)** Emission of TagRFP and mBeRFP in the 570-700 nm emission window when excited at the same power with 561 nm illumination source. (Top) Z-projection of the distal part of a leg disc showing the emission of UAS alpha-catenin-TagRFP or UAS alpha-catenin-mBeRFP. Orange and blue dashed squares point out respectively inside and outside areas used to measure the fluorescence intensity. (Bottom) Dot plot representing the fluorescence intensity in each zone. n=6 and 5 for alpha-catenin-TagRFP and alpha-catenin-mBeRFP, respectively. Black lines indicate the median. Statistical significance has been calculated using Student *t*-test. ns, not significant; *p<0.05; ***p<0.001. **E)** Five colors Z-projection of the distal part of a leg disc showing nuclei (stained with DAPI; blue), MyoII-GFP (sqh-eGFP; green), UAS alpha-catenin-mBeRFP expressed in apterous domain (yellow), E-cadherin (cyan) and actin (stained with phalloidin; red). Scale bars represent 20 µm in B, D and 10 µm in E.

We also tested the possibility of combining the mBeRFP with red-emitting fluorescent proteins or dyes (e.g., TRITC). To do that, we compared the emission rate of mBeRFP and TagRFP, both fused to alpha catenin and expressed under the control of apterous-Gal4 promoter, when excited at 561 nm and using a laser power allowing a correct visualization of TagRFP. We fixed the emission window between 570 and 700 nm to collect the maximal fluorescence rate of each protein (FigSup.2C) and we imaged leg discs expressing either alpha-catenin-TagRFP or alpha-catenin-mBeRFP. To analyze the emission of fluorescence, we compared the fluorescence intensity outside and inside the expression domain (Fig.2D). As expected, the fluorescence intensity of alpha-catenin-TagRFP is high inside the expression domain while the fluorescence intensity of alpha-catenin-mBeRFP is very close to the background intensity measured outside this area. We note however a significant residual collection of fluorescence that could be the consequence of the existence of a second excitation peak of the mBeRFP protein at 580 nm, as characterized from purified protein by Zhang group (Yang et al., 2013). To go further in this analysis, we measured the crosstalk between mBeRFP and TRITC dye after fixation of alpha-catenin-mBeRFP-expressing leg and staining for actin with phalloidin-TRITC. Similar to eGFP/mBeRFP pair, we imaged mBeRFP/TRITC pair using either spectral or classical confocal mode when the sample is illuminated with 561 nm light source. The analysis of the spectral contamination revealed an absence of crosstalk between mBeRFP and TRITC for the settings used in this assay (FigSup.2D). Thus, although mBeRFP can be weakly excited at 561 nm, its emission between 570 and 700 nm is close to the background when the laser power of 561 nm laser is low, suggesting that this fluorescent protein can be easily coupled with red-emitting fluorescent proteins without disrupting neither the observation nor the quantification.

Altogether, these data showed that mBeRFP can be combined with both green and red-emitting fluorescent proteins or dyes. In addition, this tool could even allow the detection of 5 distinct channels using 4 light sources and only 2 tracks in classical confocal mode (Fig.2E).

### mBeRFP for live imaging

The use of fluorescent proteins is mainly of interest for live imaging. We then wondered whether mBeRFP protein could be used for live imaging. To use a fluorophore in living samples, it should (1) be detected at low laser power to ensure tissue viability, (2) not be very sensitive to photobleaching. To check these characteristics, we firstly followed the adherens junctions dynamics in the apterous domain during the leg development, every 6 minutes during 4 hours (Fig.3A and Movie 1). At large scale, tissue dynamics appeared unperturbed all along the movie. At cellular scale, we are able to follow apoptotic cell extrusions from the apical surface of the epithelial leg at the expected rate (Fig.3A), based on previous observations (Monier et al., 2015). This showed that mBeRFP could be efficiently detected and used to follow dynamic processes in living samples.

**Figure3:**
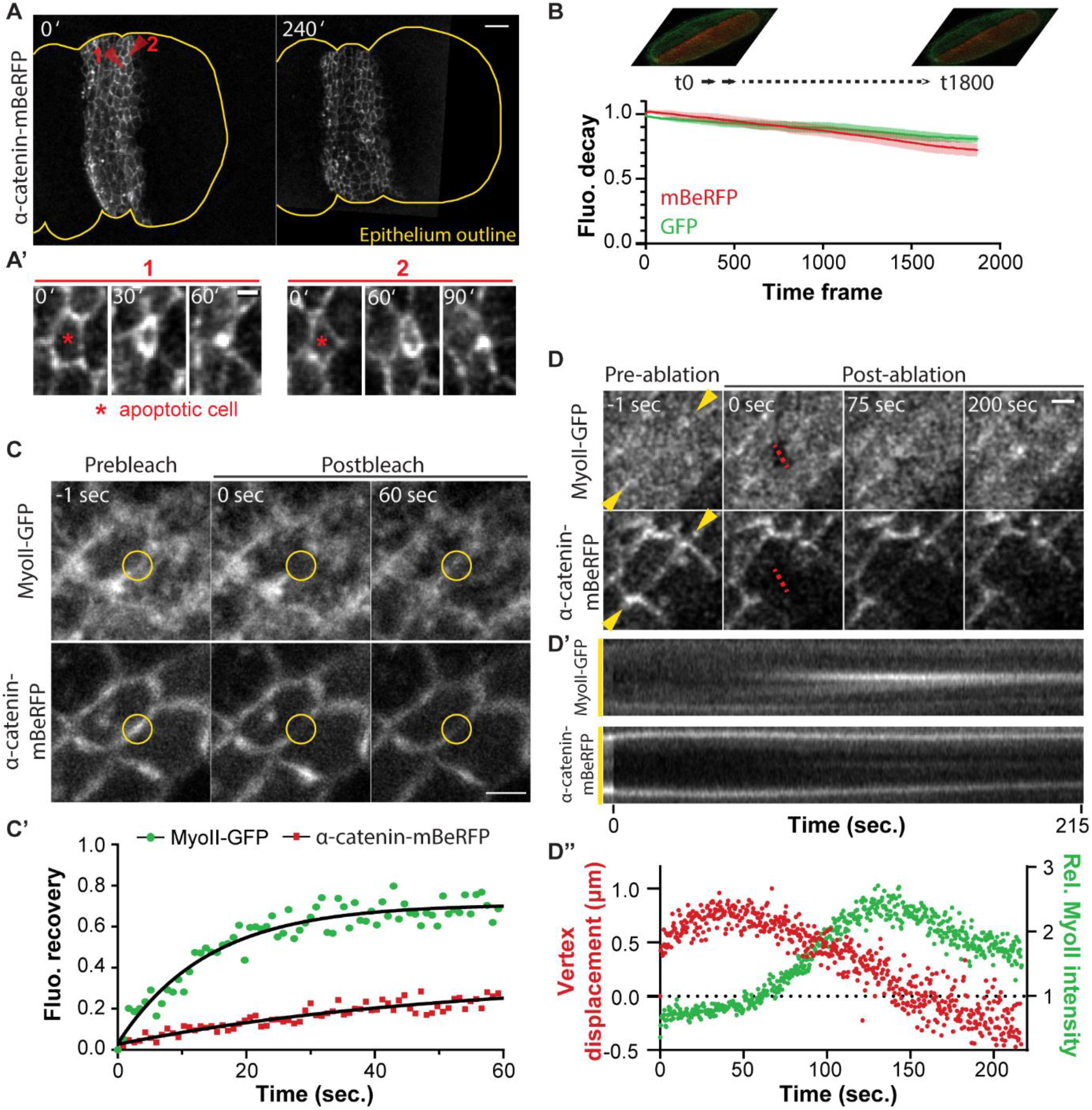
mBeRFP is a useful protein for live imaging and micromanipulation by light. **A)** Time-lapse (41-time frames) of a leg imaginal disc. A, Z-projection (41 planes) of distal part of a leg disc expressing *ap-Gal4, UAS-alpha-catenin-mBeRFP* at t0 and 4 hours after. The yellow line shows the outline of the epithelium and red arrowheads point out two apoptotic cells enlarged in A’. Scale bars represent 10 µm and 2 µm in A and A’ respectively. **B)** Fluorescence decay of mBeRFP and GFP upon 458 nm laser source excitation. A single Z-plane of wing discs expressing both MyoII-GFP (sqh-eGFP) and UAS-mBeRFP was acquired in streaming 1800 times (around 1 second / time frame). Data represented the mean of the relative fluorescence intensity +/- SEM. **C)** FRAP experiment. (Top) Confocal images showing the fluorescence recovery of alpha-catenin-mBeRFP and Myosin II-GFP after photobleaching of the area of interest (yellow circle) in *MyoII-GFP; ap-Gal4, UAS-alpha-catenin-mBeRFP* leg disc. (Bottom) Scatter plot showing the fluorescence recovery overtime of Myosin II-GFP and alpha-catenin-mBeRFP in the area of interest. Graph shows one representative experiment. Scale bar represents 2 µm. **D)** Laser ablation experiment. (Top) Confocal images showing the dynamics of adherens junctions and myosin II before and after laser ablation (red dashed line) in *MyoII-GFP; ap-Gal4, UAS-alpha-catenin-mBeRFP* leg disc. (Middle) Kymograph of adherens junctions and myosin II at the level of the yellow arrowheads. (Bottom) Scatter plots showing both the vertex displacement (red) and the fluorescence intensity of the MyoII at the level of the cut (green) overtime. Graph shows one representative experiment. Scale bar represents 2 µm.

Second, we addressed the question of the photobleaching by imaging 1800 times the same Z-plane every second, and measuring the fluorescence intensity of both GFP and mBeRFP overtime, when excited with the same 458 nm laser. This assay showed that at the end of the experiment, the fluorescence intensity of mBeRFP has dropped by 30% compared to 20% for eGFP (Fig.3B). To address this question in a more classical experiment of live imaging, we imaged a Z-stack of the wing disc every 6 minutes during 4 hours, to follow its development, and measured the fluorescence of E-cadherin-GFP and nuclear mBeRFP at each time frame (FigSup.3 and Movie 2). In this experiment, the fluorescence of both E-cadherin-GFP and nuclear mBeRFP was quite stable, thus comforting the potential practical use of this fluorescent protein for live imaging.

Finally, we reasoned that mBeRFP could constitute a very interesting tool for micromanipulation experiments and tested the possibility to use the mBeRFP fluorophore for FRAP and laser ablation. Indeed, since we can image it jointly with a GFP protein using a single excitation laser, the use of GFP/mBeRFP pair not only reduced photobleaching, but allow to image simultaneously the two fluorophores which could greatly reduce the time frame of the acquisition, a critical parameter in these types of experiments. For both experiments, we followed simultaneously MyoII-GFP and adherens junctions (α-catenin-mBeRFP) using the same excitation source. For FRAP experiments, we decided to bleach these proteins at the level of adherens junctions and followed the fluorescence recovery over time. As expected, we observed a higher stability of adherens junction protein α-catenin compared to myosin (Fig.3C) and found approximately the same rate of mobile fraction described in other systems (67% compared to 68% and 25% compared to 22%, respectively for MyoII and α-catenin (Kasza et al., 2014; Yonemura et al., 2010)). For laser ablation, in the example shown in Figure 3D, we were able to follow both the vertex displacement, based on α-catenin-mBeRFP signal, and the fluorescence intensity of the myosin at the same time. This showed that after ablation, there is a relaxation of the junction as expected, followed by a step in which the vertices come together concomitantly with a strong accumulation of myosin at the level of the cut (Fig.3D’ and D’’), suggesting the establishment of a myosin-dependent repair mechanism, similarly to what has been observed during wound repair (Antunes et al., 2013; Zulueta-Coarasa and Fernandez-Gonzalez, 2018).

## Discussion

To summarize, we have shown that mBeRFP can be easily used with low photobleaching and combined with classical fluorescent proteins, such as GFP and RFP, for multiple applications including long term live imaging and micromanipulations such as FRAP or laser ablation. Moreover, the fluorescence of this protein is not disrupted by fixation protocol (Fig.2E), allowing imaging of an additional channel in fixed tissue, and making this FP a versatile tool for imaging in model organisms. Additionally, the combination of mBeRFP with GFP allows the study of two distinct proteins of interest using the same illumination source. A single illumination source rather than two could be potentially less deleterious for tissues in terms of phototoxicity, even if this last property depends on the wavelength used as well as several other parameters (Icha et al., 2017).

If we consider excitation and emission spectra, mBeRFP could also be coupled without or with a low crosstalk with CFP, allowing to excite them together using a single light source, as previously shown (Yang et al., 2013), and YFP family proteins. Thus, it would potentially become possible to combine the most popular FRET pairs - CFP/YFP (Bajar et al., 2016) - with an additional marker coupled to mBeRFP and image them simultaneously using a single illumination source at 458 nm.

Furthermore, considering the mBeRFP excitation spectrum close to the CFP and GFP ones, we hypothesized that mBeRFP and GFP could be simultaneously observed using multiphoton microscopy, as similarly shown for CFP and GFP (Sahai et al., 2005), although it should offer better detection of each FP. In the same line, it has previously been shown that LSSmKate1, a mBeRFP sister protein, can be combined and simultaneously excited at an optimal wavelength (870 nm) with CFP and GFP for multicolor imaging using 2-photons microscopy (Piatkevich et al., 2010a).

Overall, considering the possibility to easily combine mBeRFP with other FPs, its relatively high brightness and stability in living tissues, the wide number of potential applications for live imaging as well as the possibility to use it on fixed tissues, we strongly believe that this fluorescent protein and the versatile constructs we generated will constitute important tools for cell and developmental biologists as well as biophysicists in their current research.

## Materials & Methods

### Constructs

#### pUASt-LSSmKate2, pUASt-mBeRFP and pUASt-NLSmBeRFP

LSSmKate2 and mBeRFP were individually amplified by PCR with specific primers from pLSSmKate2-N1 (Addgene #31867) and LK1-MpEF1+mBeRFP+Nos-T35S-T (gift from F. Federici, PUC, Santiago, Chile), and then cloned into the KpnI site of pUASt using InFusion technology (Clontech). The forward and reverse primers used to amplify LSSmKate2 are respectively GGCCGCGGCTCGAGGGTACCATGAGCGAGCTGATTAAGGAG and AAAGATCCTCTAGAGCTAATTAAGCTTGTGCCCCAGT. The forward primer used to amplify mBeRFP is GGCCGCGGCTCGAGGGTACCATGGTGTCTAAGGGCGA. The reverse primers used are AAAGATCCTCTAGAGTTAATTAAGTTTGTGCCCCAGT or AAAGATCCTCTAGAGTACCTTGCGCTTTTTCTTGGGAGCTCCCTCATTAAGTTTGTGCCCCAGT to generate respectively the cytoplamic or nuclear version of mBeRFP.

#### p-mBeRFP and p-NLSmBeRFP

To generate these constructs, the same strategy described above was used. Briefly, mBeRFP coding sequence was cloned into the KpnI site of phsp70, which was derived from the pUASt by removing UAS sequence.

#### pUASt--α-catenin-mBeRFP

This construct was carried out by amplifying α-catenin coding sequence from a pUAS-Dαcatenin-TagRFP (kindly provided by K. Sugimura, Kyoto University, iCeMS, Japan) then cloning in frame with mBeRFP in pUASt-mBeRFP plasmid digested with KpnI, using InFusion technology. Forward and reverse primers used to amplify α-catenin coding sequence are respectively GGCCGCGGCTCGAGGATGTTAAAACCTGATAAAATGGGCA and CCCTTAGACACCATGGCAACAGCGTCAGCAGGACT.

### Drosophila Stocks

Transgenic lines carrying UAS-LSSmKate2, UAS-mBeRFP, UAS-NLSmBeRFP and UAS-α-catenin-mBeRFP insertion at attP40 or attP2 landing site were generated in this study by standard procedures using PhiC31/attB-mediated integration. Injections were performed by the CBI Drosophila facility (Toulouse, France).

ap^md544^-Gal4 (BDSC_3041), pdm2-Gal4 (BDSC_49828), E-cad-GFP (BSCD_60584) were obtained from Bloomington Drosophila Stock Center (BDSC). sqh-eGFP^KI^[29B] (named MyoII-GFP in this article) were previously described (Ambrosini et al., 2019). Dll^EM212^-Gal4 and UAS-α-catenin-TagRFP are respectively gifts from G. Morata (CBM SO, Madrid, Spain) and K. Sugimura (Kyoto University, iCeMS, Japan).

### Immunofluorescence

Imaginal leg and wing discs were respectively dissected 2 hours after pupae formation or at third instar larval stage in PBS 1X. Tissues were fixed in 4% paraformaldéhyde (PFA 4%) for 20’, then washed in PBS and mounted in Vectashield containing or not either DAPI or phalloidin rhodamine (Vectors laboratories). For Figure 2E, samples were washed in PBS-Triton 0.3%-BSA 1% (BBT) after fixation and incubated overnight at 4°C with rat anti-E-cadherin antibody (Developmental Studies Hybridoma Bank – DCAD2) diluted at 1:50 in BBT. After washes in BBT, tissues were incubated with 1:200 anti-rat IgG 647 for 2 h at room temperature with 1:500 phalloidin TRITC (Fisher Scientific). Then, samples were washed in PBS-Triton 0.3% and mounted on slides in Vectashield with DAPI (Vectors laboratories).

For the analysis of the photobleaching over time (Fig.3B), samples were washed in PBS directly after fixation in PFA, re-suspended in Scheider medium and mounted on a slide to mimic live experiments and avoid tissue motion.

### Drosophila live samples

For live imaging, FRAP and laser ablation experiments (Fig.3A, C-D), imaginal leg disc from ap-Gal4>UAS-NLSmBeRFP or sqh-eGFP^KI^[29B]; ap-Gal4>UAS-mBeRFP animals were dissected from prepupae in Schneider medium supplemented with 15% fetal calf serum, 0.5% penicillin-streptomycin and 2 μg/mL 20-hydroxyecdysone (Sigma-Aldrich, H5142) and mounted on slides.

For photobleaching experiments (FigSup.3), imaginal wing disc from sqh-RFPt^KI^[3B]; pdm2-Gal4>UAS-NLSmBeRFP animals were dissected from early third-instar larvae in Schneider’s insect medium supplemented with 15% fetal calf serum, 0.5% penicillin-streptomycin and mounted on slides.

### Confocal, Spectral and RIM microscopy

Images were acquired with a Zeiss LSM880 confocal microscope (Carl Zeiss) equipped with a 30 mW 405 nm diode laser, a 34,4 mW 458/488/514 nm argon multiline laser, a 13 mW 561 nm DPSS laser, and a 3 mW 633 nm HeNe laser on a Zeiss Axio Observer microscope. Microscope was also fitted with a Plan-Apochromat 40x/NA 1.3 Oil DIC UV-IR M27, 63X C-Apochromat NA 1.2 Water Corr (Fig.3C-D FRAP and laser ablation) and 40X C-Apochromat NA 1.2 Water Corr (Fig.3A-B live imaging and photobleaching assay) objectives. Z-stacks were acquired using either the laser scanning confocal mode or the spectral mode and the adapted illumination source. A linear unmixing process was done following spectral acquisition to separate each fluorophore based on their spectra.

Random Illumination Microscopy was then performed using a home-made system previously described (Mangeat et al., 2021). Images were acquired every 4 ms using an inverted microscope (TEi Nikon) equipped with a 100x magnification, 1.49 N.A. objective (CFI SR APO 100XH ON 1.49 NIKON) and Abbelight two sCMOS camera (ORCA-fusion, Hamamatsu) system band pass filters (Semrock): FF01-514/30-25 for GFP, FF01-630/92-25 for mBeRFP, Fast diode lasers (Oxxius) with wavelength centered at 445 nm (LBX 445 100 CSB OE) were used for the excitation of both GFP and mBeRFP. A spatial light phase binary modulator (QXGA fourth dimensions) was conjugated to the image plane to create speckle random illumination. Image reconstruction was then performed as previously detailed (Mangeat et al., 2021) with the upgraded version of AlgoRIM V1.2 : https://github.com/teamRIM/tutoRIM.

### FRAP experiments

Fluorescence Recovery After Photobleaching (FRAP) was performed on the LSM880 confocal microscope described above. For FRAP experiments, GFP and mBeRFP were excited simultaneously using the 458 nm laser source and photons were collected by GaAsP detectors. Photobleaching was set through the bleaching module provided on ZEN software. Briefly, the region of interest was illuminated 5 times using a bleach dwell time of 0.60 μs/pixel with 100 % 488 nm laser power. Images were acquired every 900 ms, 10 times before bleaching and 100 times post bleaching.

### Laser ablation

Laser ablation experiments were carried out using a pulsed DPSS laser (532 nm, pulse length 1.5 ns, repetition rate up to 1 kHz, 3.5 μJ/pulse) steered by a galvanometer-based laser scanning device (DPSS-532 and UGA-42, from Rapp OptoElectronic, Hamburg, Germany) and mounted on a LSM880 confocal microscope (Carl Zeiss). The microscope was fitted with a 63x C-Apochromat NA 1.2 Water Corr objective (Carl Zeiss). Photo-ablation of the apical junction was done in the focal plane by illuminating at 95-98 % laser power during 500 ms. Time-lapse was acquired using a 458 nm light source, every 370 ms, during 5 s before and at least 200 s after ablation, with a pixel size of 0.13 μm/pixel, and photons were collected in two emission windows (from 490 nm to 550 nm for eGFP and from 600 nm to 700 nm for mBeRFP).

### Quantification

To assess the crosstalk, the photobleaching or the fluorescence intensity, the mean of fluorescence intensity was measured in the region of interest using ImageJ (Fig.2B-D, 3B-D, FigSup.1B, 2D, 3).

The crosstalk in classical laser scanning mode between GFP and mBeRFP was calculated as the difference between the rate of eGFP emitted in the mBeRFP emission window when imaged at 458 nm with the spectral mode and the corresponding rate in the classical mode. The crosstalk in classical laser scanning mode between mBeRFP and TRITC was calculated as the difference between the rate of mBeRFP emitted in the TRITC emission window when imaged at 561 nm with the spectral mode and the corresponding rate in the classical mode.

Vertex displacement was evaluated on kymograph using a homemade macro in ImageJ.

### Statistical analysis

The normality of data sets was determined using Prism 8 (Graph Pad). A Student *t*-test was used to assess the significance in Fig.2D and FigSup.1B.

## Acknowledgements

We would like to thank Bruno Monier for his constructive comments on the manuscript and Thomas Mangeat, from the LITC platform, for his help on RIM experiments.

## Author contributions

E.M. conceived and performed the experiments, analyzed and quantified the data and wrote the paper. M.S. supervised the project, participated in writing the paper, and provided the funding.

## References

Ambrosini, A., Rayer, M., Monier, B. and Suzanne, M. (2019). Mechanical Function of the Nucleus in Force Generation during Epithelial Morphogenesis. Dev Cell 50, 197-211.e195.

Antunes, M., Pereira, T., Cordeiro, J. V., Almeida, L. and Jacinto, A. (2013). Coordinated waves of actomyosin flow and apical cell constriction immediately after wounding. J Cell Biol 202, 365–379.

Bajar, B. T., Wang, E. S., Zhang, S., Lin, M. Z. and Chu, J. (2016). A Guide to Fluorescent Protein FRET Pairs. Sensors (Basel) 16.

Boulina, M., Samarajeewa, H., Baker, J. D., Kim, M. D. and Chiba, A. (2013). Live imaging of multicolor-labeled cells in Drosophila. Development 140, 1605–1613.

Chalfie, M., Tu, Y., Euskirchen, G., Ward, W. W. and Prasher, D. C. (1994). Green fluorescent protein as a marker for gene expression. Science 263, 802–805.

Chudakov, D. M., Matz, M. V., Lukyanov, S. and Lukyanov, K. A. (2010). Fluorescent proteins and their applications in imaging living cells and tissues. Physiol Rev 90, 1103–1163.

Gorgidze, L. A., Oshemkova, S. A. and Vorobjev, I. A. (1998). Blue light inhibits mitosis in tissue culture cells. Biosci Rep 18, 215–224.

Heppert, J. K., Dickinson, D. J., Pani, A. M., Higgins, C. D., Steward, A., Ahringer, J., Kuhn, J. R. and Goldstein, B. (2016). Comparative assessment of fluorescent proteins for in vivo imaging in an animal model system. Mol Biol Cell 27, 3385–3394.

Icha, J., Weber, M., Waters, J. C. and Norden, C. (2017). Phototoxicity in live fluorescence microscopy, and how to avoid it. Bioessays 39.

Kasza, K. E., Farrell, D. L. and Zallen, J. A. (2014). Spatiotemporal control of epithelial remodeling by regulated myosin phosphorylation. Proc Natl Acad Sci U S A 111, 11732–11737.

Kogure, T., Karasawa, S., Araki, T., Saito, K., Kinjo, M. and Miyawaki, A. (2006). A fluorescent variant of a protein from the stony coral Montipora facilitates dual-color single-laser fluorescence cross-correlation spectroscopy. Nat Biotechnol 24, 577–581.

Mangeat, T., Labouesse, S., Allain, M., Negash, A., Martin, E., Guenole, A., Poincloux, R., Estibal, C., Bouissou, A., Cantaloube, S., et al. (2021). Super-resolved live-cell imaging using Random Illumination Microscopy. Cell Reports Methods 1.

Monier, B., Gettings, M., Gay, G., Mangeat, T., Schott, S., Guarner, A. and Suzanne, M. (2015). Apico-basal forces exerted by apoptotic cells drive epithelium folding. Nature 518, 245–248.

Piatkevich, K. D., Hulit, J., Subach, O. M., Wu, B., Abdulla, A., Segall, J. E. and Verkhusha, V. V. (2010a). Monomeric red fluorescent proteins with a large Stokes shift. Proc Natl Acad Sci U S A 107, 5369–5374.

Piatkevich, K. D., Malashkevich, V. N., Almo, S. C. and Verkhusha, V. V. (2010b). Engineering ESPT pathways based on structural analysis of LSSmKate red fluorescent proteins with large Stokes shift. J Am Chem Soc 132, 10762–10770.

Sahai, E., Wyckoff, J., Philippar, U., Segall, J. E., Gertler, F. and Condeelis, J. (2005). Simultaneous imaging of GFP, CFP and collagen in tumors in vivo using multiphoton microscopy. BMC Biotechnol 5, 14.

Shimomura, O., Johnson, F. H. and Saiga, Y. (1962). Extraction, purification and properties of aequorin, a bioluminescent protein from the luminous hydromedusan, Aequorea. J Cell Comp Physiol 59, 223–239.

Subach, O. M., Cranfill, P. J., Davidson, M. W. and Verkhusha, V. V. (2011). An enhanced monomeric blue fluorescent protein with the high chemical stability of the chromophore. PLoS One 6, e28674.

Thorn, K. (2017). Genetically encoded fluorescent tags. Mol Biol Cell 28, 848–857.

Wiedenmann, J., Oswald, F. and Nienhaus, G. U. (2009). Fluorescent proteins for live cell imaging: opportunities, limitations, and challenges. IUBMB Life 61, 1029–1042.

Yang, J., Wang, L., Yang, F., Luo, H., Xu, L., Lu, J., Zeng, S. and Zhang, Z. (2013). mBeRFP, an improved large stokes shift red fluorescent protein. PLoS One 8, e64849.

Yonemura, S., Wada, Y., Watanabe, T., Nagafuchi, A. and Shibata, M. (2010). alpha-Catenin as a tension transducer that induces adherens junction development. Nat Cell Biol 12, 533–542.

Zimmermann, T., Rietdorf, J. and Pepperkok, R. (2003). Spectral imaging and its applications in live cell microscopy. FEBS Lett 546, 87–92.

Zulueta-Coarasa, T. and Fernandez-Gonzalez, R. (2018). Dynamic force patterns promote collective cell movements during embryonic wound repair. Nature Physics 14, 750–758.

